# A multimodal computational pipeline for 3D histology of the human brain

**DOI:** 10.1101/2020.02.10.941948

**Authors:** Matteo Mancini, Adrià Casamitjana, Loic Peter, Eleanor Robinson, Shauna Crampsie, David L. Thomas, Janice L. Holton, Zane Jaunmuktane, Juan Eugenio Iglesias

**Author notes:** For correspondence (MM), (JEI).

## Abstract

*Ex vivo* imaging enables analysis of the human brain at a level of detail that is not possible *in vivo* with MRI. In particular, histology can be used to study brain tissue at the microscopic level, using a wide array of different stains that highlight different microanatomical features. Complementing MRI with histology has important applications in *ex vivo* atlas building and in modeling the link between microstructure and macroscopic MR signal. However, histology requires sectioning tissue, hence distorting its 3D structure, particularly in larger human samples. Here, we present an open-source computational pipeline to produce 3D consistent histology reconstructions of the human brain. The pipeline relies on a volumetric MRI scan that serves as undistorted reference, and on an intermediate imaging modality (blockface photography) that bridges the gap between MRI and histology. We present results on 3D histology reconstruction of a whole human hemisphere.

## 1 Introduction

### 1.1 Motivation

One of the major challenges in the quest to understand the human brain as a complex system is characterizing its multiscale organization. From a macroscopic anatomical perspective, the brain is subdivided into distinct structures and presents well-defined landmarks, for instance its gyri and sulci. At a much finer scale, neurons are interconnected to form complex circuits. These circuits give rise to features at coarser scales, e.g., the laminar organization of the cortex [1]. Although these distinct levels of organization may seem separate, they are, in fact, deeply linked and mutually influential. One clear example is given by Alzheimer’s disease, which can be described by the interactions between proteinopathy at the microscale and distributed, network-level disruptions at the macroscale [2]. Understanding these multi-scale mechanisms requires a multi-scale map of the brain. However, the current concept of brain mapping is closely linked to the specific tool used to construct cartographic representations, and thus to the spatial scale of the tool. As a result, the different organizational principles can only be observed in a scale-specific fashion with dedicated tools [3].

Multi-scale imaging of the human brain is therefore necessarily multimodal, as different modalities are needed to study different scales. While a macroscopic anatomical picture of the brain is easily acquired non invasively or *ex vivo* with magnetic resonance imaging (MRI), finer characterization requires histological procedures and microscopy. Histology and MRI are highly complementary modalities: histology produces excellent contrast at the microscopic scale using dedicated stains that target different microanatomical or cytoarchitectural features, but it is a 2D modality that also inevitably introduces distortions in the tissue during blocking and sectioning. MRI does not yield microscopic resolution, but produces undistorted 3D volumes. Therefore, the combination of these two modalities offers a solution to the problem of imaging the human brain at high resolution in 3D. In fact, successful large-scale projects like BigBrain [4] or the Allen Atlas [5] have shown important advancements in terms of creating new whole-brain atlases with cellular-level resolution. Creating such atlases requires spatial alignment of images (“registration”) at the macroscopic (MRI) and microscopic (histology) scales. Such registration produces 3D-consistent histological volumes, and is often called “3D histology reconstruction” [6].

Despite remarkable efforts like BigBrain, 3D histology reconstruction still presents obstacles: it requires manual intervention and tailored equipment, which leads to poor scalability and limited applicability in other experimental and clinical studies. Specifically, three main issues can be identified:

1. Cutting and sectioning of tissue for histology introduces distortions (stretching, tearing, folding, cracking, see [6]) that are specific to each section. Therefore, an external volumetric reference (typically an MRI scan) is required to produce an unbiased registration. Without such a reference, naïve pairwise registration leads to accumulation of errors (“z-shift”) and spurious straightening of curved structures (the so-called “banana effect”, [7]).
2. Large differences in contrast and resolution between MRI and histology, combined with potential inhomogeneous staining and the aforementioned sectioning artifacts, make the alignment of these two modalities a difficult inter-modality registration problem.
3. The large size of the human brain, compared with most animal models, requires cutting the tissue in blocks for whole-brain analyses, with the resulting need to reassemble the blocks [6], which is a part-to-whole registration problem [8, 9]. This problem can be solved with whole brain microtomes, although such microtomes are only available in few selected sites around the world. Moreover, they exacerbate the sectioning artifacts, due to the much larger surface area of the sections.

Even when a reference MRI is available, 3D histology reconstruction is an ill-posed problem, as errors in the – typically linear – registration between the MRI and the histological stack can also be explained by nonlinearly deforming the histological sections. To overcome this problem, a common strategy is to use an intermediate modality for registration purposes. Several works [10–13] have used photographs of the block during sectioning (so-called blockface photos). While these photos do not have nearly as much contrast or resolution as the histology, they have the advantage of being free from sectioning artifacts. Therefore, they can be corrected for illumination and perspective and then stacked into blockface volumes, which constitutes a useful stepping stone between histology and MRI.

Other potential ways of obtaining 3D consistent volumes at microscopic scale include optical coherence tomography (OCT, [14]), polarized light imaging (PLI, [15]) and cleared tissue microscopy [16]. Notably, large-scale microtomes are not a feasible solution with these techniques, as they are inherently limited to small tissue blocks. In fact, despite technological advances in terms of both increasingly larger samples [17–19] and novel microscopic acquisitions [20, 21], complete multi-scale characterization of the human brain as a whole organ still requires the use of complementary tools able to cover all the biologically relevant scales. This inherent limitation makes the development of inter-modality workflows a necessity, especially as cutting-edge research starts to target the whole human body [22].

### 1.2 Related work

There is a growing literature on the topic of combining histology with other modalities, encompassing different scopes and subdomains in medical imaging beyond neuroimaging. Here we will provide a brief survey of the approaches proposed so far for this multimodal problem, presenting first the ones focused on specific samples or small organs, and then addressing methods targeting whole organs.

Despite not being spatially comprehensive, approaches based on selected regions of interest (ROIs) have a high clinical relevance, mainly because of their potential applications in oncology. It is no surprise, then, that several studies have combined MRI with histopathology for cancer applications: examples include breast cancer [23, 24], pancreatic tumors [25] and gliomas [26]. These examples lay the foundation for future 3D histopathology, especially given the parallel effort in tridimensional reconstruction for confocal microscopy [27].

In human neuroimaging, combined MRI-histology also holds great potential because of the cross-scale nature of neurological diseases. Recent studies have proposed ROI-based approaches to better understand the microscopic substrate of pathologies such as amyotrophic lateral sclerosis [28], epilepsy [13] and Alzheimer’s disease [29]. The main targets mostly include subcortical structures, including thalamus [30, 31], hippocampus [32, 33], nucleus accumbens [34], and pedunculopontine nucleus [35], among others [10]. In addition to the practical advantage of dealing with well-defined structures, the focus on the subcortex is due mainly to its implications in neurological diseases and psychiatric disorders.

Related approaches in terms of target size include methods focused on small organs. Several studies have proposed combined MRI-histology approaches for the prostate [36, 37], lymphoid structures [38], mammary glands [39], and kidneys [40]. Another comparable application is the study of small animal brains, in particular rodents [41–43] and small monkeys [44,45]. A common element to most of the pipelines mentioned so far is the histological section stacking procedure, with registration of consecutive sections as the central step, usually using a combination of rigid and non-linear transformations, and taking advantages of application-specific landmarks where available. To further facilitate the registration process, an interesting approach recently proposed by several works is based on the use of 3D printing to create personalized molds on the basis of MRI data [37, 46–48], introducing shape constraints for the subsequent cutting procedure.

In contrast to the large body of existing work in 3D reconstruction of small samples, the literature on whole-brain approaches is rather limited. Apart from the major initiatives already mentioned above [4, 5], to the best of our knowledge there are only two other studies that have targeted either the entire brain [49] or a major portion of the cerebrum [50]. These studies relied on the availability of whole brain microtomes, allowing to build on a section stacking procedure similar to sample-targeted approaches. As already mentioned, this is a significant limitation for the scalability of tridimensional histology, and also strongly limits attempts to leverage on new microscale technologies: most of the new advancements require small samples and therefore cutting the brain in blocks.

### 1.3 Contribution

As explained above, the number of potential approaches to probe microstructure in small samples is increasing, but directly adopting such techniques in whole human brain histology-MRI reconstruction is infeasible. Therefore, the ability to reconstruct a whole brain distribution of a given microscopic biomarker from a set of smaller samples is crucial to build multimodal, multi-scale maps of the human brain. In this article, we present an open-source computational pipeline to reconstruct human brain volumes from histological sections with *ex vivo* MRI, blockface photographs and a standard microtome. To the best of our knowledge, this is the first approach able to reconstruct whole-brain 3D histology from a block-based cutting protocol. Since a highly specialized whole hemisphere microtome is not required, the proposed pipeline can be used by any research site with access to a standard microtome and an MRI scanner. The pipeline relies on a number of 2D and 3D image registration methods, some standard, and some developed specifically for this pipeline, which have already been introduced at conferences [51, 52]. Here we introduce the pipeline as a whole and present results on the reconstruction of a whole human hemisphere.

## 2 Materials and Methods

In this section, we describe the pipeline for 3D histology reconstruction, including the data acquisition protocol and computational processing, as summarized in figure 1.

**Figure 1.**
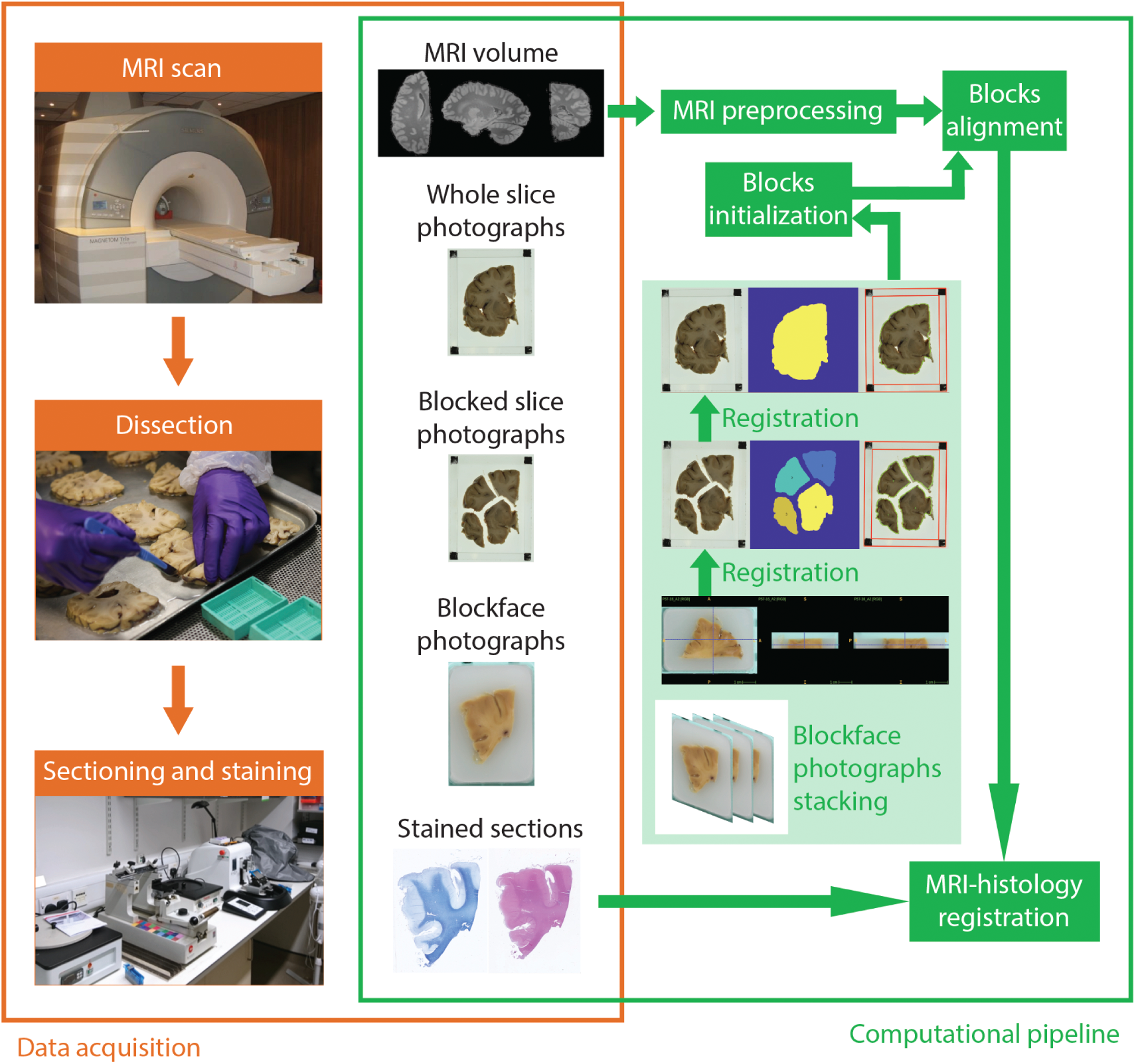
Workflow of data acquisition (orange) and computational processing (green. The *ex vivo* brain is scanned, dissected, sectioned and stained, providing data for the pipeline: the MRI volume, the whole and blocked slice photographs (dissection), the blockface photographs (sectioning), and the stained sections. The flowchart illustrates the main steps of the pipeline: stacking of blockface photographs; registration of blockface volumes to slice photographs; blocks initialization; block mosaic-preprocessed MRI alignment; and MRI-histology registration.

### 2.1 Data acquisition

#### 2.1.1 Specimen preparation and *ex vivo* MRI scanning

In this study, we use tissue donated for research to the Queen Square Brain Bank for Neurological Disorders (QSBB). The brain donation program and protocols have received ethical approval for research by the NRES Committee London - Central and tissue is stored for research under a license issued by the Human Tissue Authority (No. 12198). According to the standard protocol at QSBB, fresh brains are first hemisected. The right hemisphere is frozen, while the left one is fixed in 10% neutral buffered formalin.

For *ex vivo* MRI scanning, it is important to immerse the brain in a fluid, in order to avoid susceptibility artifacts at tissue-air interfaces around the edges of the brain. Using the fixative as a medium for this purpose is problematic due to the high proton density of formalin, which quickly saturates the MR signal, thus greatly reducing the dynamic range of the acquired images. Instead, we use Fluorinert (perfluorocarbon), a proton-free fluid which matches the magnetic susceptibility of brain tissue but has no MR signal, so it is invisible in MR images. Immersion in Fluorinert yields excellent *ex vivo* contrast, and it is known not to affect subsequent histological analysis of the tissue for a wide array of stains, for up to a week of immersion [53].

MRI data are acquired on a 3T Siemens MAGNETOM Prisma scanner. T1-weighted MR imaging is a common choice *in vivo* due to its excellent contrast between gray and white matter. However, death and fixation induce a cross linking of proteins that greatly shortens T1 relaxation times of brain tissue, reducing T1 contrast *ex vivo*. Instead, we use a T2-weighted sequence (optimised long echo train 3D fast spin echo, [54]) with parameters: TR=500 ms, TEeff=69 ms, BW=558 Hz*/*Px, echo spacing=4.96 ms, echo train length=58, 10 averages, with 400 µm isotropic resolution. The hemisphere is scanned in a container filled with Fluorinert, with the medial surface facing up. We place a 3D printed hollow box between the specimen and the container lid in order to ensure full immersion in the fluid. Sample slices of the acquisition are shown in figure 2*a*.

**Figure 2.**
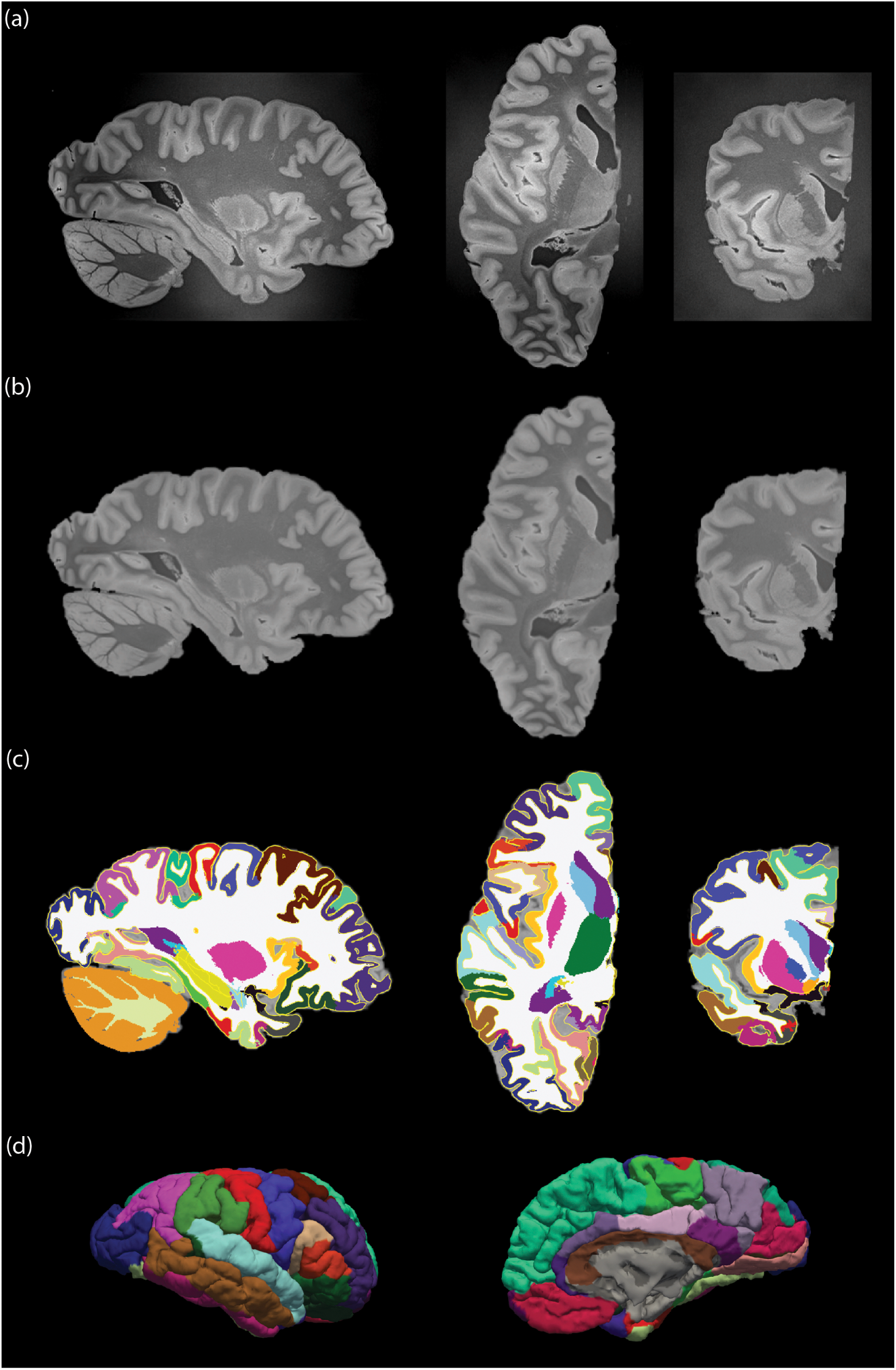
Overview of the main MRI preprocessing steps. (a) The raw *ex vivo* MRI data as acquired, showed in sagittal (left), axial (middle) and coronal (right) views; (b) brain extraction and bias field correction given by SAMSEG; (c) segmentation or subcortical structures given by SAMSEG, combined with cortical segmentation and parcellation provided by FreeSurfer, with the outline of the reconstructed surfaces in yellow; (d) 3D rendering of the cortical surface from lateral (left) and medical (right) views. The color coding of the segmentation and parcellation follows the FreeSurfer convention.

#### 2.1.2 Specimen dissection and slice photography

After MRI scanning, the hemisphere is dissected following the procedure illustrated in figure 3. First, the brainstem is detached with a transection perpendicular to its axis below the mammillary body (figure 3*c-d*), and the cerebellum is separated from the cerebrum. The three structures are then dissected independently. The cerebrum is first cut into 10 mm-thick coronal slices, starting from the mammillary body and proceeding in both anterior and posterior directions (figure 3*e-f*). In a similar way, the cerebellum and the brainstem are sliced in sagittal and axial orientation, respectively, with the same thickness. All the slices are then photographed on both sides (posterior and anterior for the cerebrum, rostral and caudal for the brainstem, medial and lateral for the cerebellum) inside a rectangular frame of known dimensions (internal boundary: 120 mm × 90 mm), and thickness equal to the slice thickness (10 mm). The frame enables perspective correction and pixel size calibration in subsequent steps of the pipeline. We will refer to these images as “whole slice photographs”, which will be useful to initialize the 3D histology reconstruction.

**Figure 3.**
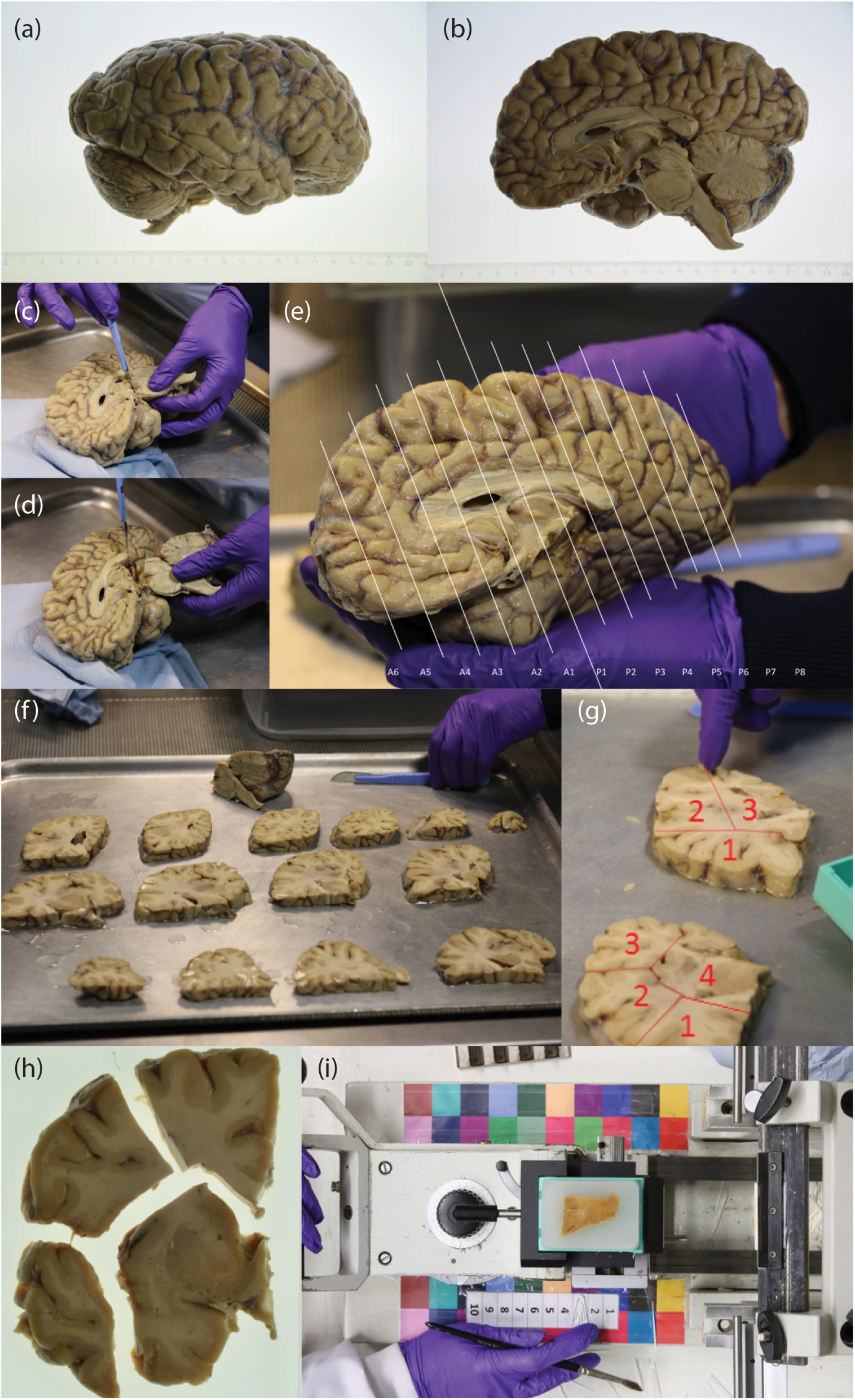
Overview of the dissection procedure. (a-b) Brain hemisphere from lateral (a) and medial (b) views; (c) incision; (d) subsequent brainstem transection below the mamillary body; (e) overview of how the anterior and posterior slices are cut; (f) cerebrum slices and remaining brainstem and cerebellum portions; (g) example of block cut planning; (h) example of blocked slice photograph; (i) example of blockface photograph, including the microtome setup.

The cerebrum sections are further cut into blocks that fit into 74×52 mm cassettes, seeking to minimize the number of blocks while trying to preserve the integrity of subcortical structures (figure 3*g*). In our datasets, cutting the brainstem and the cerebellum in blocks was never necessary, since they always fit directly into the cassettes. Photographs of the blocked slices, where the blocks were slightly pulled apart to clearly expose their boundaries, were taken using the same frame as for the whole slices, both from the anterior and posterior side. We will refer to these images as “blocked slice photographs”.

All slice photographs (whole and blocked) are taken with a Nikon D5100 camera mounted over the samples. Consistent image contrast across samples is ensured by manually setting: ISO=100; one-shot auto focus using a single point in the center of the image (which is aligned with the center of the slice); and f/20 aperture for large depth of field. The shutter speed was computed automatically by the camera to compensate for variations in lighting level. Examples of slice photographs are shown in figure 4*a-b*.

**Figure 4.**
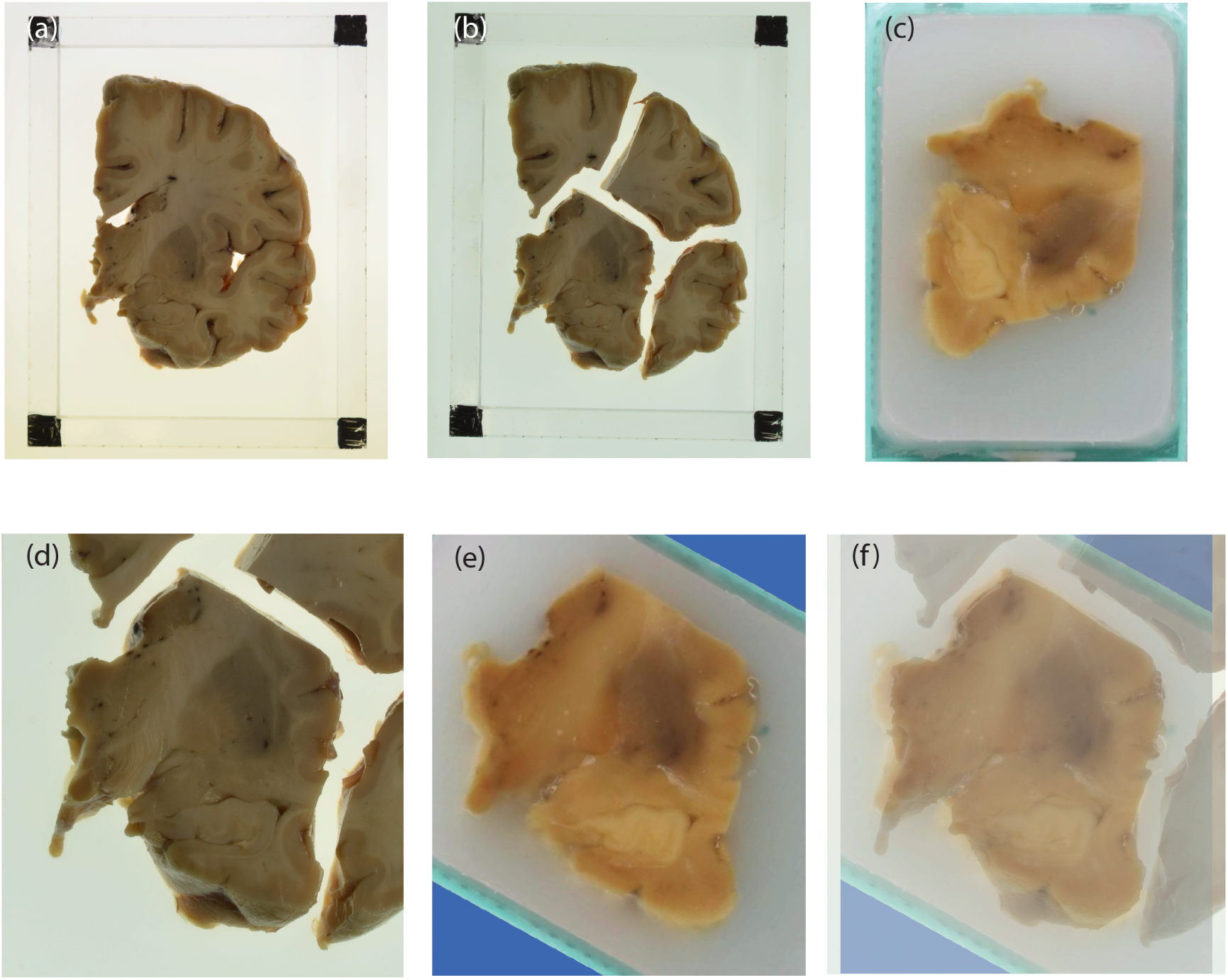
An example of blocked slice photograph to blockface photograph registration. (a) A sample whole slice photograph; (b) the related blocked slice photograph; (c) corresponding blockface photograph of the bottom left block; (d) block of interest in the blocked slice photograph; (e) aligned blockface photograph; (f) blocked slice photograph with blockface photograph overlaid in transparency.

#### 2.1.3 Tissue processing, sectioning and blockface photography

All blocks are processed for paraffin wax embedding, and subsequently sectioned with a sledge microtome at 25 µm thickness. Before cutting each section, a photograph is taken with a camera mounted above the microtome, set in a fixed position that is approximately perpendicular to the slicing plane (figure 3*i*). We will refer to these images as “blockface photographs”. Since these photographs will need to be perspective corrected, pixel size calibrated, and co-registered (since keeping the camera absolutely still is not possible), we printed and glued two checkerboard patterns with maximally distinct colors [55] to the microtome, which facilitates subsequent registration. The photographs are taken at 24MPx resolution with a Canon EOS 750D camera. As for the slice photographs, we use manual settings to ensure consistency of image appearance across sections: ISO=200; white balance = fluorescent light source; one-shot auto focus using a single point in the center of the image (which is aligned with the center of the tissue block); and crucially, a narrow aperture (f/13) for large depth of field, thus ensuring sharpness of objects not exactly in focus. An example of blockface photograph is shown in figure 4*c*.

#### 2.1.4 Staining and digitization

Tissue blocks are classified into two groups: “interesting” (those including subcortical structures in the cerebrum, the medial blocks in the cerebellum and all the blocks in the brainstem) and “uninteresting” (all other blocks). For interesting blocks, we mount on glass slides and stain two consecutive sections every 10 (i.e., every 250 µm) with two routine histological stains: hematoxylin and eosin (H&E) and Luxol Fast Blue (LFB). For uninteresting blocks, the frequency is one every 20 instead (i.e., every 500 µm). The sections are mounted on 75×50 mm glass slides. We also mount 2 additional slides every 10 (interesting blocks) or 200 sections (uninteresting), unstained, for potential future use.

Stained sections are digitized with a flatbed scanner (Epson Perfection V850) using its transparency mode at 6,400 DPI (i.e., 3.97 µm resolution). This resolution is sufficient for 3D histology reconstruction purposes. Selected sections are also digitized at microscopic resolution (40×) using an Olympus VS120 microscope / slide scanner, and linearly aligned to the corresponding images acquired with the flatbed scanner using NiftyReg [56].

### 2.2 Computational pipeline

After completing the acquisition of MRI and histological data, the following pipeline is used to compute the 3D reconstruction.

#### 2.2.1 Ex vivo MRI

The T2-weighted MRI scan is preprocessed using a Bayesian segmentation algorithm (SAMSEG, [57]) that simultaneously registers, segments and bias field corrects the scan (figure 2*b*). To reflect the presence of just the left hemisphere in the images, we modified the SAMSEG probabilistic atlas by manually setting to 1 the probability of background (and to 0 for all other classes) for all voxels in the right half of the atlas. Our modified SAMSEG produces a bias field corrected scan, as well as segmentations for 22 brain structures: cerebral white matter, cerebellum white matter, brainstem, ventral diencephalon, optic chiasm, cerebral cortex, cerebellum cortex, caudate, hippocampus, amygdala, accumbens area, lateral ventricle, inferior lateral ventricle, 3^*rd*^ ventricle, 4^*th*^ ventricle, 5^*th*^ ventricle, cerebrospinal fluid, vessel, choroid plexus, thalamus, putamen, and pallidum.

After SAMSEG, we used FreeSurfer [58] to extract and parcellate the cortical ribbon. Specifically, we used the SAMSEG cerebral white matter segmentation and followed these steps (figure 2*c-d*): *(i)* extraction of a triangular mesh from the cerebral white matter segmentation with marching cubes [59]; *(ii)* inflation of the mesh and mapping to spherical coordinates [60]; *(iii)* topology correction [61]; *(iv)* reconstruction of white matter and pial surfaces [62]; and *(v)* cortical parcellation [63].

### 2.2.2 Blockface photographs

In an ideal scenario, the blockface photographs would be perfectly aligned without any need for processing. However, the position of the arm holding the camera can suffer from small perturbations due to vibrations in the furniture and walls, operation of the microtome, and other external factors. Therefore, it is necessary to align the photographs before further processing. For this purpose, we first create a global reference image (“microtome reference”), which is a photograph of the microtome with an empty cassette. On this image, we manually mark the four corners of the cassette, and delineate two masks: one over the checkerboard patterns, and another over a band around the edges of the cassette.

We use the microtome reference to perspective correct and calibrate the pixel size of all other blockface photographs, by propagating the location of the four corners of the cassette. For this purpose, we first select the photograph half way through the block, which we will refer to as “block reference”. The microtome reference is registered to the block reference to propagate the location of the cassette corners as follows: *(i)* we compute salient points and SURF features [64] on both images, and discard those outside the checkerboard in the microtome reference; *(ii)* we match the salient points; *(iii)* we use random sample consensus (RANSAC, [65]) to fit an homography (perspective) transform; and *(iv)* we refine the registration to accurately align the cassettes, by repeating the procedure in steps *(i-iii)*, but with two differences: we use the mask for the cassette edges instead of the checkerboard mask, and we add an Euclidean distance term to the matching, since the cassettes are already in coarse alignment. We use RANSAC to make the registration robust against different positions of the microtome handle and appearance of the tissue block. The final transform is used to propagate to the block reference the location of the manually labeled cassette corners, as well as the mask for the checkerboard patterns.

Once we have estimated the checkerboard mask and cassette corners for the blockface reference, we register all photographs in the block to the reference using steps *(i-iii)* of the procedure described above. Finally, we compute an homography transform between the four cassette corners and coordinates (1, 1), (1, 740), (520, 1), (520, 740), and use it to resample the blockface photographs into 520×740 pixel images with known pixel size equal to 100 µm, where the corners of the image coincide with the corners of the cassette. These images can be safely stacked into a single volume, with *z* resolution equal to 25 µm. We note that the in-plane pixel size is slightly overestimated due to the fact that the actual blockface is slightly closer to the camera than the cassette. Moreover, the *z* resolution is also corrupted by inaccuracies in the section thickness provided by the microtome. Nevertheless, these voxel size errors do not represent a problem in practice because tissue shrinks during processing, and both the pixel size and section thickness need to be corrected in subsequent steps of the computational pipeline anyway.

The blockface photograph module is completed by a supervised segmentation algorithm, which discriminates tissue versus background wax. We use a fully convolutional network (FCN, [66]) trained on manual segmentations made on 50 randomly selected (perspective corrected, pixel size calibrated) images from different blocks. The FCN was built by on top of the VGG16 network [67], with preinitialized weights for transfer learning. While the FCN operates in 2D, stacking the automated segmentations yields a 3D mask that is spatially smooth, due to the 3D smoothness of the underlying images.

#### 2.2.3 Slice photographs

The slice photographs are crucial to initialize the registration of the tissue blocks. Processing of both whole and blocked slice photographs begins by segmenting the corners of the frame, which are painted in black, using a FCN similar to the one used for blockface photographs. This time we used 20 manually labeled images for training, which is enough, given the simplicity of the problem. The center of gravity of the four largest clusters are identified as the corners. Then, four sets of parallel lines are fit to the gradient magnitude images, to identify the internal and external boundaries of the frame. The internal corners are computed as the intersections of the internal boundaries, and used to fit an homography to correct for perspective and calibrate the pixel size to 100 µm (i.e., 1,200×900 pixels), in a similar way as for the blockface photographs.

Next, the blocked slice photographs (perspective and pixel size calibrated) are segmented into foreground and background using a simple Gaussian mixture model (GMM) with two components, optimized with the Expectation Maximization (EM) algorithm [68]. The resulting mask is overlaid onto the corresponding image and displayed on a simple graphical user interface (GUI), where a user assigns block numbers to the different connected components of the binary mask, producing a multilabel segmentation of the different blocks.

#### 2.2.4 Digitized histological sections

Processing of digitized histological sections has two components: segmentation and intensity standardization. For segmentation, we used two FCNs, one for LFB and one for H&E, trained on 50 randomly selected sections each. Simple intensity standardization was carried out using only the pixels inside the masks, by matching their histogram to the average histogram of the foreground pixels of the training dataset.

#### 2.2.5 Linear alignment of blocks

The advantage of using blockface photography as intermediate modality is that the registration to the volumetric reference (in our case, the MRI volume) is approximately linear. In our pipeline, we first use the slice and blockface photographs to initialize the registration between blockface volumes and the MRI. Then, we use a dedicated joint registration algorithm to optimize the alignment.

To initialize the registration, we start by calculating three different sets of 2D linear transforms. First, we use SURF and RANSAC (see section ”Blockface photographs”) to rigidly register whole slice photographs of the lateral / anterior / inferior face of each block to the medial / posterior / superior face of the neighboring lateral / anterior / inferior block. These images are nearly identical, so registration is easy and accurate. Second, we use SURF/RANSAC to rigidly align each block in the blocked slice photographs (using the available multi-label masks) to the corresponding whole slice photograph. Despite being a whole-to-part registration problem, SURF/RANSAC produces accurate solutions, since the photographs are of the same objects and acquired in the same illumination conditions. And third, we estimate a similarity transform (i.e., translation, rotation and scaling) between each block in the blocked slice photographs and the approximately corresponding image in the blockface photograph volume, using mutual information as cost function [69–71]. The target blockface photograph is the first one where the whole block is completely visible, which we estimate by finding the first section in which the ratio between surface area and its maximum across the block is at least 2*/*3. The surface areas are estimated with the masks produced by the FCN. An example of the alignment between a blocked slice photograph and the related blockface one is showed in figure 4*d-f*.

Given these sets of 2D transforms, initializing the blocks in the cerebellum, cerebrum and brainstem is straightforward. For the cerebrum, a reference slice is first chosen (the one corresponding to the mammillary bodies, in our case). Then, for each block, one simply concatenates its corresponding blocked-slice-to-whole-slice transform, along with all the whole-slice-to-whole-slice transforms between the slice at hand and the reference. Finally, a shift in the anterior-posterior direction is computed by each block, which is simply equal to the slice thickness (10 mm) multiplied by the (signed) number of slices between the slice at hand and the reference. The cerebellum and brainstem are processed the same way, with two differences: the blocked-slice-to-whole-slice transform is not needed (since slices are not blocked), and the shifts occur in the medial-lateral (cerebellum) and inferior-superior directions (brainstem). Finally, the three sets of blocks (cerebrum, cerebellum and brainstem) are manually aligned to the preprocessed reference MRI using a rigid transform.

Once the transforms for each block have been initialized, they are optimized using a joint hierarchical registration algorithm. The details of the methods can be found in [52], but we summarize them here for completeness. Each blockface volume has an associated spatial transform that has a set of 8 parameters, corresponding to 3D translation (3 parameters), 3D rotation (3), in-plane scaling (1), and scaling along the thickness direction (1). The cost function of the registration combines a data term and two regularizers. The former is simply the correlation of edge maps. The first regularizer is a customized penalty term that encourages the sum of the soft deformed masks to be equal to the binary mask of the MRI. This regularizer penalizes gaps between blocks (the sum is zero, whereas the target is 1) as well as overlapping blocks (sum is 2, target is 1), and also encourages the surface of the whole hemisphere to be the same for the MRI and the mosaic of blockface volumes. The second regularizer penalizes deviations of the global scaling of each block (i.e., the cubic root of the determinant of its linear transformation matrix) from an empirical value, which represents the expected tissue shrinkage, and which we derive from the blocked-slice-to-blockface registrations.

The optimization procedure is hierarchical, in order to exploit prior knowledge on the cutting procedure. There are five levels of hierarchy. At the first level, blocks are grouped into three sets (cerebrum, cerebellum, brainstem), each of which undergoes an independent rigid registration. At the second level, the cerebrum blocks are grouped into corresponding slices and can only rotate or translate simultaneously in the slice plane. At the third level, individual translation and rotation are allowed, with the addition of a scaling factor that is common to all blocks. At the fourth level, each block is allowed its own scaling factor. At the fifth and final level, each block has its own transform. We use the L-BFGS [72] algorithm for numerical optimization of the cost function.

#### 2.2.6 Registration with histology

Once the blockface volumes have been linearly registered to the reference MRI volume, it is straightforward to resample the MRI onto the plane of any of the blockface photographs. Since the correspondence between blockface photographs and digitized stained sections is known and deterministic, the MRI can thus be resampled on the planes corresponding to these sections. Furthermore, the 2D resampled MRI slices can be masked with the automated segmentations of the blockface photographs provided by the FCN. This process yields a stack of resampled, segmented MR images that have direct correspondence to the images in the histology stack.

To 3D reconstruct the histology for each of the two stains (LFB and H&E), we used a method that we presented in [51], and which we summarize here for completeness. The goal is to register a stack of histological sections to a corresponding stack of resampled 2D MRI slices. First, the histology stack is put into coarse alignment by linearly registering each section to the corresponding MR image. For this purpose, we used the linear registration module in NiftyReg [56], which relies on a block matching approach and mutual information. Then, we compute a set of nonlinear registrations parameterized by stationary velocity fields (SVF, [73]), as implemented in NiftyReg. Using SVFs has three advantages. First, the corresponding deformations are guaranteed to be diffeomorphic and thus invertible; second, inversion is achieved simply by changing the sign of the SVF; and third, composition of transforms can be approximated by the sum of the corresponding SVFs. The set of registrations includes: *(i)* inter-modality registrations between each histological section and the corresponding MRI slice, computed with mutual information; *(ii)* intra-modality registrations, between each histological section and its two nearest neighbors in the stack, computed with local normalized cross-correlation; and *(iii)* intra-modality registrations, between each MRI slice and its two nearest neighbors in the stack, also computed with local normalized cross-correlation.

While one could use the inter-modality registrations – i.e., subset *(i)* – directly to obtain a 3D reconstruction, this approach is known to produce volumes that are jagged, due to inconsistencies in the registrations of neighboring image pairs. At the opposite end of the spectrum, an alternative approach is to use only the intra-modality registrations for histology, i.e., subset *(ii)*, but this method leads to accumulation of errors along the stack (“z-shift”) and straightening of curved structures (“banana effect”, see [6]). Instead, our method [51] achieves the best of both approaches by combining all three subsets. The method relies on a spanning tree of unknown, “true” deformations connecting all the images in the two stacks. Then, all the registrations in the three subsets can be seen as noisy measurements of compositions of transforms in the set of true deformations. Within this model, Bayesian inference is used to compute the most likely set of underlying true deformations that gave rise to the observed data. This approach produces 3D reconstructions that are both smooth and robust against z-shift and banana effect. The intuition behind it is that subset *(i)* aligns the two stacks; subset *(ii)* ensures the smoothness, and subset *(iii)* enables us to undo the banana effect incurred by subset *(ii)*.

After running all the modules of the computational pipeline, correspondence between spatial locations in the MRI and digitized histological sections can be obtained by concatenating 3D (blockface volume to MRI) and 2D spatial transforms (histology to resampled MRI).

## 3 Results

A left human hemisphere was processed with this pipeline. In this section we present intermediate and final results, including 3D tissue blocks, alignment with MRI and 3D histology reconstruction.

### 3.1 Building blockface volumes

figure 5 displays results from the first fundamental step of our 3D histology pipeline: assembling blockface photographs into tridimensional volumes. The reconstructed views of a sample cerebral block (axial and sagittal, figure 5*a*) present a smooth and consistent profile, demonstrating how effective the co-registration strategy we adopted is.

**Figure 5.**
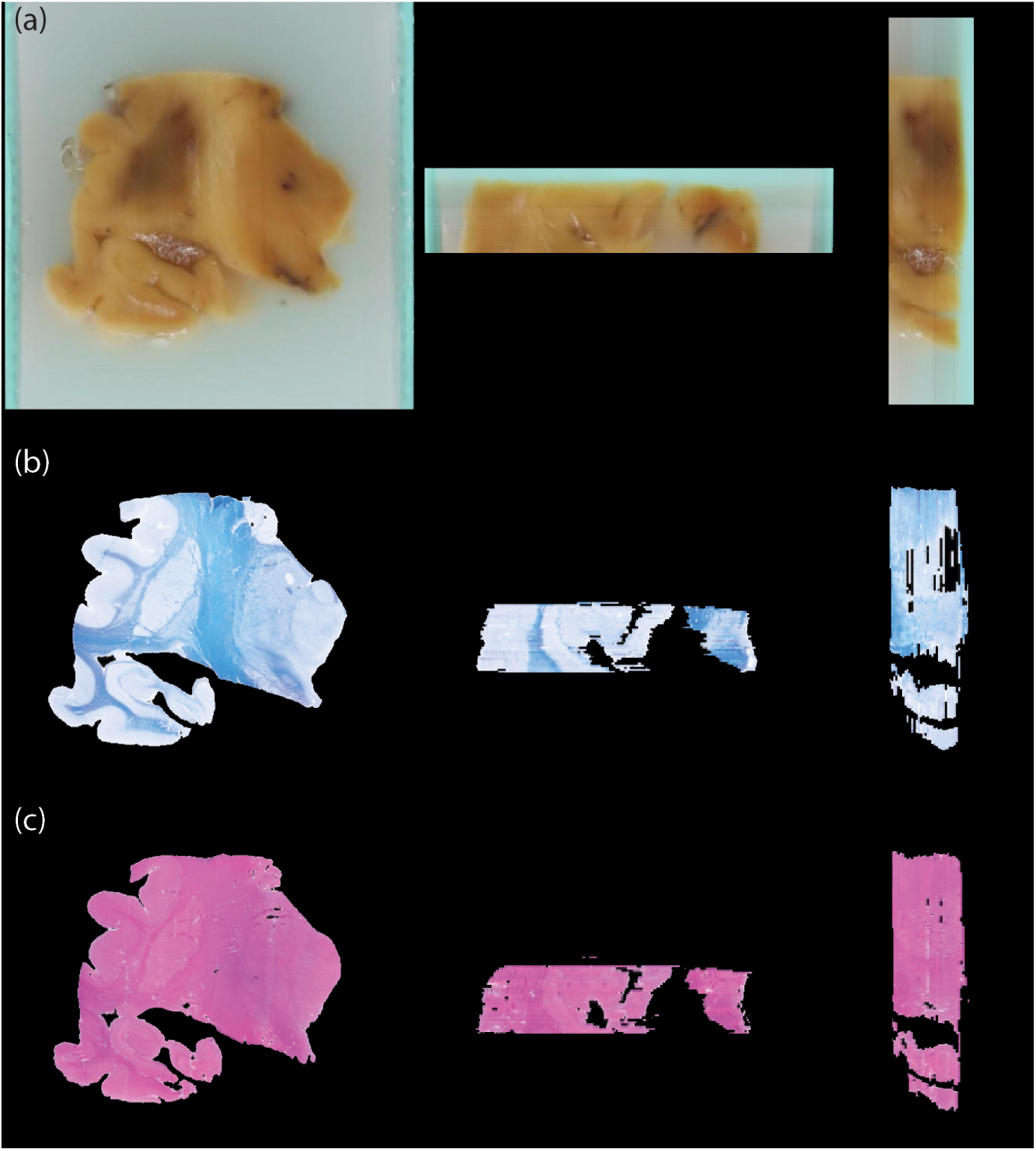
Blockface and histological volumes. Example of reconstructed blockface volumes (a) from three different views (coronal on the left, axial in the centre, and sagittal on the right). The same views are also showed for the correspondent histological volumes, stained with LFB (b) and H&E (c).

figure 5*b-c* provides a first glimpse of how the blockface volumes are used as a stepping stone in the overall process of fusing MRI and histology. The figure shows the histological sections (LFB and H&E) corresponding to the blockface photograph in figure 5*a*. For easier visualization, we show the sections after nonrigid registration with the MRI, which indirectly aligns them with the blockface photograph as well. Therefore, the volumes are consistent with each other – but they also shows some differences because of inevitable artifacts that occur during sectioning.

### 3.2 Effective alignment of MRI reference and tissue blocks

Once all the blockface volumes have been assembled and initialized, a tridimensional mosaic representing the brain is obtained (figure 6*a-b*). Notably, the overall shape and the coarse subdivisions (cerebrum, cerebellum, brainstem) are a good first approximation, although misalignment in the gyrification patterns and in the antero-posterior organization are clearly noticeable. When compared with the actual MRI reference (whose whole brain mask is surface rendered in transparent green), there is a clear mismatch in their shapes.

**Figure 6.**
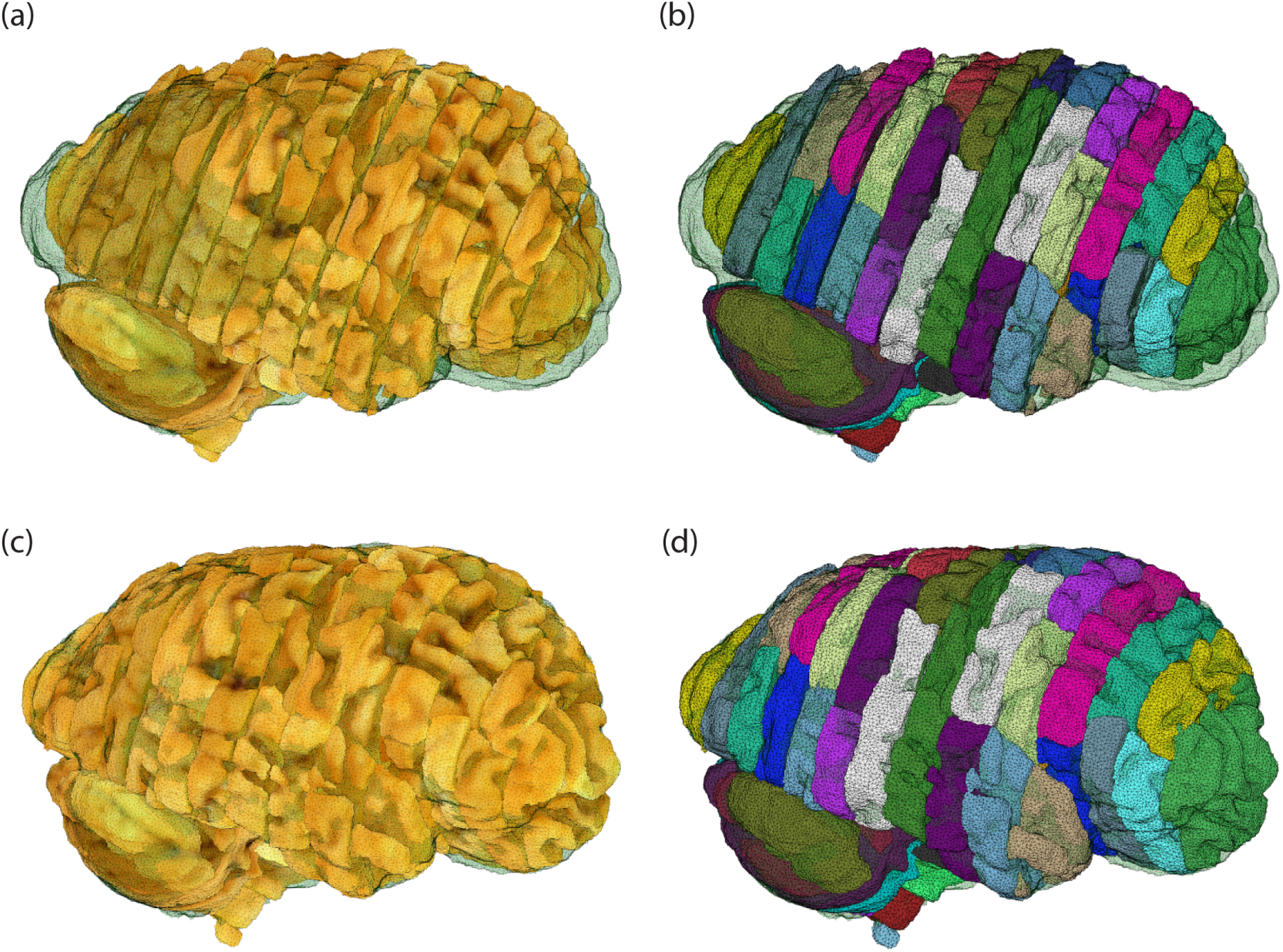
Initialization and refined blockface mosaics. A surface rendering of the whole brain mask derived from the MRI is overlaid in green. (a) Blockface volume for the whole hemisphere resulting from the initialization with the slice photographs; the color for rendering is taken from the blockface photographs. (b) Same volume as (a), where each block is rendered in a different color. (c,d) Same volumes as (a,b) after joint registration.

Our joint registration procedure yields the refined mosaic shown in figure 6*c-d*: not only the blocks match the MRI reference well, but also yield a more consistent and smoother gyrification pattern. ***Supplementary video 1*** thoroughly illustrates the joint registration procedure, and gives a more detailed overview of both the initialized and the refined mosaics – highlighting how the consistency of brain structures improves as a result of the spatial alignment.

### 3.3 Navigating histological sections in 3D

Navigating through different scales of co-registered histology-MRI data, e.g., to localize and examine a section of interest, is one of the most immediate applications of this pipeline. Given the estimated transformations between blockface photographs and histological sections, it is possible to use the refined, MRI-aligned blockface mosaic to jointly explore the histological sections and the MRI in a common three-dimensional space. figure 7 shows the same block as figure 5, this time overlaid on the MRI volume. The figure displays the alignment of the different brain structures across MRI and histology, both in the sectioning (coronal) and reconstructed planes (axial, sagittal). Moreover, it also highlights the smoothness of the reconstructed histology.

**Figure 7.**
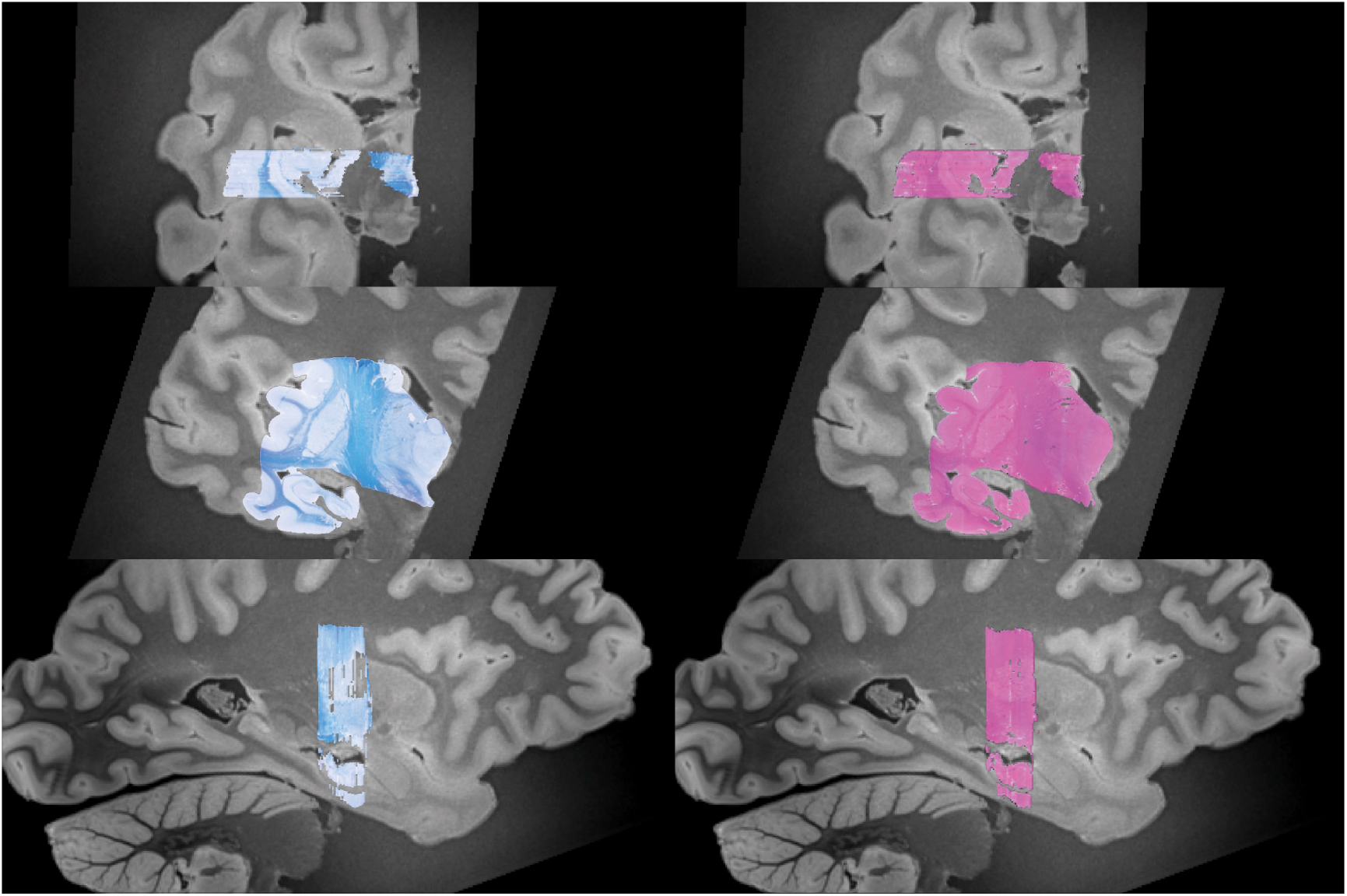
MRI-histology alignment. Histological sections from a sample block aligned with the MRI volume as a result of the refinement algorithm, showed from three different views (top: axial; middle: coronal; bottom: sagittal) and for the two stains (left: LFB; right: H&E).

figure 8*a* shows a more detailed example of navigation and alignment, displaying an LFB section on the corresponding resampled MRI plane. The red square delineates a 5×5 mm patch magnifying the basal ganglia (figure 8*b-d*), specifically the boundaries between the putamen and the two segments of the globus pallidus. The alignment is excellent, and the histology reveals details that are effectively invisible in MRI (even at sub-millimeter resolution). Likewise, the green square marks a 30×30 mm patch magnifying hippocampal head and the amygdala (figure 8*e-g*). Once again, the alignment between the boundaries across the two modalities is qualitatively very high.

**Figure 8.**
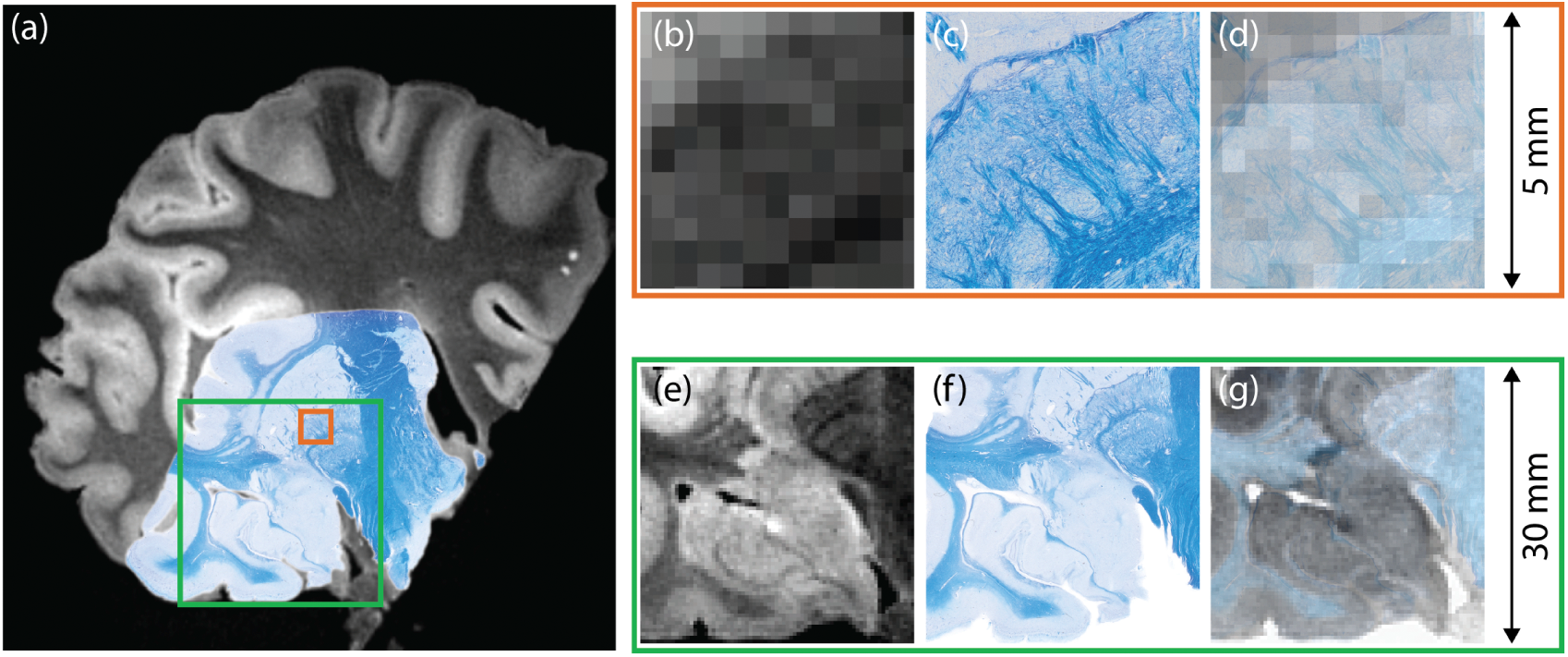
Registration between MRI and histological sections. (a) Sample MRI slice with registered LFB section overlaid; (b) magnified view of the 5×5 mm area marked in red; (c) Corresponding region on LFB section; (d) MRI and LFB overlapped in transparency; (e-g) magnification of the 30×30 mm area marked with the green square in (a).

Navigation is further exemplified in ***Supplementary video 2***, which combines the multi-modal imaging data with the segmentations from SAMSEG and FreeSurfer. Starting from the brain surface, the perspective slowly focuses on the basal ganglia, highlighting the alignment between blockface photograph, MRI and histology across different scales – from centimeters to microns per pixel. Finally, it is also possible to reconstruct and entire histology volume by mosaicking all the sections in 3D. An example at 0.4 mm resolution (i.e., the voxel size of the MRI) is showed in figure 9.

**Figure 9.**
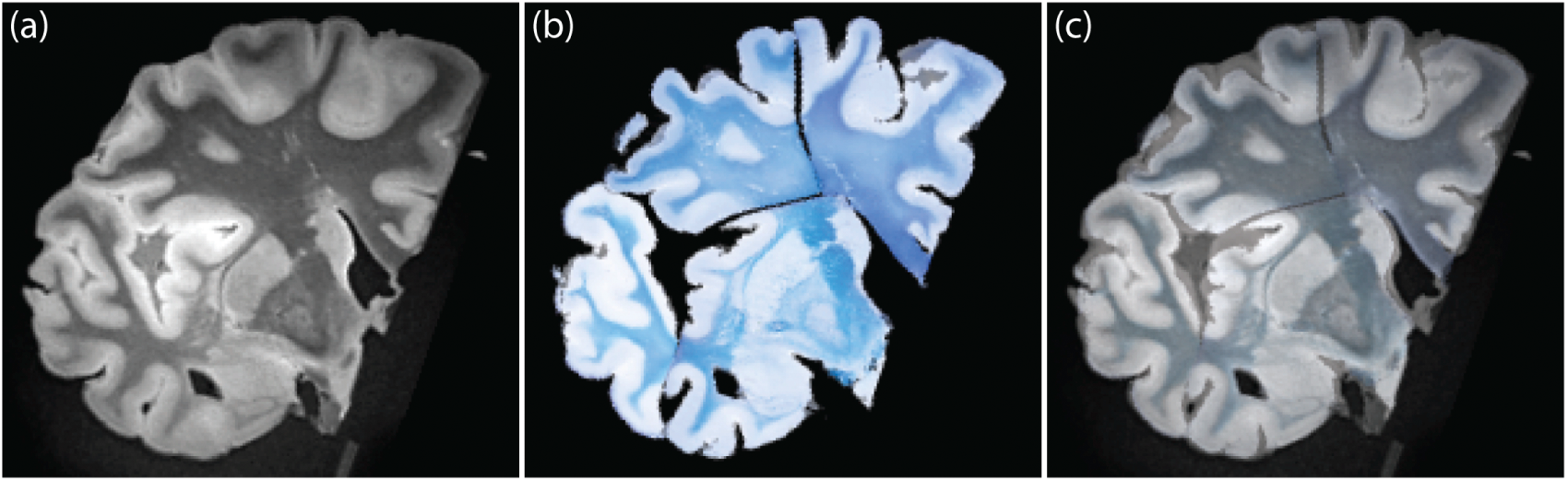
Example of whole brain reconstruction. (a) A sample coronal MRI slice; (b) corresponding slice in reconstructed LFB mosaic; (c) overlay of (a) and (b) in transparency.

## 4 Discussion

In this paper, we have described a scalable and reproducible method for MRI-informed 3D histology. This includes both a protocol for tissue cutting and processing, without any requirements apart from a standard microtome, and a computational pipeline that requires minimal manual intervention. We have also presented results for a single hemisphere of a human brain processed with the pipeline. As the reader may have observed, the proposed procedure consists of several steps, each with different design choices. In this section, we want to provide further context for such choices and discuss how they influenced our results.

### 4.1 MRI as a reference for histology

The transition towards 3D histology is becoming necessary for a more complete study of biological specimens at a microscopic level. Such specimens are tridimensional objects and therefore a 3D characterization is required. Unfortunately, histology requires by definition the loss of tridimensional shape for the target object, and without prior information on the original shape, it is not possible to reconstruct a 3D volume in a way that is coherent with the original object.

As outlined in the introduction, MRI is a powerful tool when imaging entire organs, especially for large human organs such as the brain. There are virtually no comparable alternatives when one considers also how different acquisition sequences can be used to obtain a diverse collection of contrasts. Unfortunately, the wide spatial coverage comes at the expense of a coarser resolution. Although the idea of combining MRI and histology to achieve the best of both worlds is reasonable, the practical implementation is not straightforward. For instance, registration of a single histological section leads to a difficult slice-to-volume problem [8], with the disadvantages of high sensitivity to initialization conditions and challenging multi-modal registration issues [6]. This is why our approach relies on a volume-based approach, by first assembling together histological sections from the same block into volumes. The overall registration problem still requires further steps as discussed in the following paragraphs.

### 4.2 Choice of intermediate modality

An important choice in designing a pipeline for tridimensional histology regards the intermediate modality, which will serve as a stepping stone between the stained sections and the MRI data. In this paper, we relied on blockface photographs. Since they keep structural information before the cutting procedure [6], they allow on one hand to assume a linear relationship with the MRI reference, and on the other hand to deterministically map histological sections and blockface photographs. However, there are other potential candidates for this role, most notably OCT, PLI and clearing techniques.

OCT is an interferometry technique based on near-infrared light [14]. OCT volumes can be acquired during the sectioning procedure of a sample, with the important difference (compared with histology) that the data are acquired *before* sectioning. Therefore, geometric distortion is avoided and direct stacking of the 2D images yields a 3D consistent volume. While OCT produces images with excellent contrast at high resolution, it is much more costly to acquire than photographs, especially in terms of time [74, 75]: imaging a single 20 mm thick human tissue block at 5×5×50um takes several days of uninterrupted data acquisition.

Also based on optical imaging, PLI exploits the transmission of polarized light to give a quantitative estimate of fiber orientation and inclination angles for a given point in a tissue section [15]. This technique has great potential to study white matter at the mesoscopic and microscopic scales [76], but still requires sectioning.

Tissue clearing [16] is a powerful solution that avoids sectioning. Clearing methods can make opaque tissue transparent and, combined with fluorescent labeling tools, they offer a new way of probing microscopic structures [77, 78]. As a consequence of the transparency, there is no need to slice the sample and it is sufficient to adjust the focus of the microscope on the plane of interest. The main drawbacks of this technique are the long time needed to process the tissue, the size limits for clearable samples and the limited depth of antibody penetration in the cleared tissue.

It is evident then that the choice of an intermediate modality is a trade-off between result quality and costs. Therefore, we chose to use blockface photographic imaging as it is cheap and fast, and as a result ideal for larger scale studies. The poorer contrast of blockface photographs compared to the alternatives is indeed a limit of the approach presented here, but in the context of the whole pipeline, it serves its purpose well as intermediate modality.

### 4.3 Assumptions of registration-based pipelines

In our pipeline, we have made the assumption that the deformation between the MRI and the tissue blocks (imaged via photography) is linear. This is an approximation: even with fixed tissue, cutting into slices and blocks introduces small nonlinear deformations, particularly near the cut boundaries. Moreover, tissue processing also introduces nonlinear distortions, even if minimal.

Another potential error source for the registration is the imperfection of the automated segmentation of the blockface photographs, particularly when the tissue is concave: since paraffin is not opaque, the apparent surface of the tissue on the blockface is often overestimated. As these automatically generated masks are crucial in the regularization of our linear registration, their oversegmentation may have directly affected the quality of our linear alignment and thus our results. This problem could be mitigated by integrating the linear and nonlinear registration algorithms: given the nonlinear registration of the histological sections, we could take advantage of the superior contrast of the stained sections and their more accurate automated segmentations, in order to improve the linear registration. In a similar fashion, the newly improved linear alignment could be used to refine the non-linear registration, and so on. Future work will explore this direction.

### 4.4 Improving non-linear registration

The final step of this pipeline and the last transformation needed for MRI-histology alignment is given by the registration between the resampled MRI slices and the related histological sections. In this case, the non-linearities are considerable, and the cross-modality nature of the problem requires the use of inter-modality metrics, such as mutual information. As it has been previously shown [79], approaches based on mutual information often perform poorly in inter-modality registration, and therefore represent a bottleneck when registering MRI images and stained sections – even when they are already initialized with our joint linear registration method. In order to improve the alignment, an important direction to explore is the use of synthesis, i.e. estimating MRI contrast from the histology (or vice versa) to reduce the registration to an easier intra-modality problem. Recent advances with architectures based on generative adversarial networks have shown great potential for this specific problem [80, 81], even with specific applications to medical imaging [82].

### 4.5 Future applications

In this paper we have presented, as a preliminary practical application of our pipeline, the chance of exploring histological sections through the related MRI volume. This is only the tip of the ice for the potential applications of our approach. In the overarching strategy of our current project, the next step is to acquire a larger set of brains, manually segment the stained sections, and exploit 3D histology to create a probabilistic atlas of the human brain at the subregion level.

As opposed to existing techniques, our pipeline will be able to build whole-brain datasets from stained sections, allowing us to build atlases that are much more detailed than current templates. Moreover, since specific staining agents and immunohistochemistry techniques can enhance different microscopic details, the approach described here opens the door to draw new multimodal maps of the human brain [83], in a scalable and reproducible way. With the advancement of microscopy-oriented techniques (e.g., [77,84–88]), we believe that closing the gap with macroscopic modalities is crucial.

Another potential application is the use of histology for development of MRI-based biomarkers and quantitative imaging ([89]): as *ex vivo* validation with histology becomes more common, the neuroimaging field will largely benefit from 3D histology.

Finally, aligned MRI and histology also have the potential to lead to ultra-large-scale microscopy: so far it is possible to create a large dataset with several images acquired using electron microscopy, by stitching them together to cover the entire histological section [90]. In this perspective, our approach could potentially lead to the ability to navigate a brain volume and then retrieve ultra-high-resolution details from a point of interest. This would be the new frontier of multi-scale imaging and would of course create new challenges, since the associated storage and processing requirements are highly demanding.

### 4.6 Conclusion

We have presented a pipeline to effectively obtain high-resolution 3D images of the human brain using histology and MRI. The related code and the acquired data are publicly available ^1^, and we plan to use the presented methods to build a high-resolution computational atlas of the human brain based on 3D reconstructed histology. As increasingly more advanced macroscale and microscale techniques to study the brain become available, the open-source tools we have presented here will have a key role in bridging together the two ends of the scale.

## 5 Acknowledgments

This work was primarily supported by a Starting Grant from the European Research Council (ERC), awarded to JEI (grant agreement 677697, project “BUNGEE-TOOLS”). DLT was supported by the UCL Leonard Wolfson Experimental Neurology Centre (PR/ylr/18575). MM is funded by the Wellcome Trust through a Sir Henry Wellcome Postdoctoral Fellowship. JLH is supported by the Multiple System Atrophy Trust; the Multiple System Atrophy Coalition; Fund Sophia, managed by the King Baudouin Foundation and Karin & Sten Mortstedt CBD Solutions. Queen Square Brain Bank is supported by the Reta Lila Weston Institute for Neurological Studies and the Medical Research Council UK. This research was supported by the National Institute for Health Research University College London Hospitals Biomedical Research Centre. The authors would like to thank Efthymios Maneas for his help with the plexiglass frame, and Danaiil Nikitichev for his help 3D printing the box to help keep the samples immersed in Fluorinert during MRI scanning.

## Footnotes

1 We will make our code repository public upon publication of this manuscript.

## References

1. Nicola Palomero-Gallagher and Karl Zilles. Cortical layers: Cyto-, myelo-, receptor-and synaptic architecture in human cortical areas. Neuroimage, 2017.

2. David T Jones, David S Knopman, Jeffrey L Gunter, Jonathan Graff-Radford, Prashanthi Vemuri, Bradley F Boeve, Ronald C Petersen, Michael W Weiner, and Clifford R Jack Jr. Cascading network failure across the alzheimer’s disease spectrum. Brain, 139(2):547–562, 2015.

3. Katrin Amunts and Karl Zilles. Architectonic mapping of the human brain beyond brodmann. Neuron, 88(6):1086–1107, 2015.

4. Katrin Amunts, Claude Lepage, Louis Borgeat, Hartmut Mohlberg, Timo Dickscheid, Marc-Étienne Rousseau, Sebastian Bludau, Pierre-Louis Bazin, Lindsay B Lewis, Ana-Maria Oros-Peusquens, et al. Bigbrain: an ultrahigh-resolution 3d human brain model. Science, 340(6139):1472–1475, 2013.

5. Song-Lin Ding, Joshua J Royall, Susan M Sunkin, Lydia Ng, Benjamin AC Facer, Phil Lesnar, Angie Guillozet-Bongaarts, Bergen McMurray, Aaron Szafer, Tim A Dolbeare, et al. Comprehensive cellular-resolution atlas of the adult human brain. Journal of Comparative Neurology, 524(16):3127–3481, 2016.

6. Jonas Pichat, Juan Eugenio Iglesias, Tarek Yousry, Sébastien Ourselin, and Marc Modat. A survey of methods for 3d histology reconstruction. Medical image analysis, 46:73–105, 2018.

7. Grégoire Malandain, Eric Bardinet, Koen Nelissen, and Wim Vanduffel. Fusion of autoradiographs with an mr volume using 2-d and 3-d linear transformations. NeuroImage, 23(1):111–127, 2004.

8. Enzo Ferrante and Nikos Paragios. Slice-to-volume medical image registration: A survey. Medical image analysis, 39:101–123, 2017.

9. Jonas Pichat, Eugenio Iglesias, Sotiris Nousias, Tarek Yousry, Sébastien Ourselin, and Marc Modat. Part-to-whole registration of histology and mri using shape elements. In Proceedings of the IEEE International Conference on Computer Vision, pages 107–115, 2017.

10. Eduardo Joaquim Lopes Alho, Ana Tereza Di Lorenzo Alho, Lea Grinberg, Edson Amaro, Gláucia Aparecida Bento Dos Santos, Rafael Emídio da Silva, Ricardo Caires Neves, Maryana Alegro, Daniel Boari Coelho, Manoel Jacobsen Teixeira, et al. High thickness histological sections as alternative to study the three-dimensional microscopic human sub-cortical neuroanatomy. Brain Structure and Function, 223(3):1121–1132, 2018.

11. Jessica Lebenberg, A-S Hérard, Albertine Dubois, Julien Dauguet, Vincent Frouin, Marc Dhenain, Philippe Hantraye, and Thierry Delzescaux. Validation of mri-based 3d digital atlas registration with histological and autoradiographic volumes: an anatomofunctional transgenic mouse brain imaging study. Neuroimage, 51(3):1037–1046, 2010.

12. Ann S Choe, Yurui Gao, Xia Li, Keegan B Compton, Iwona Stepniewska, and Adam W Anderson. Accuracy of image registration between mri and light microscopy in the ex vivo brain. Magnetic resonance imaging, 29(5):683–692, 2011.

13. Jacopo Annese, DM Sforza, M Dubach, D Bowden, and Arthur W Toga. Postmortem high-resolution 3-dimensional imaging of the primate brain: blockface imaging of perfusion stained tissue. Neuroimage, 30(1):61–69, 2006.

14. David Huang, Eric A Swanson, Charles P Lin, Joel S Schuman, William G Stinson, Warren Chang, Michael R Hee, Thomas Flotte, Kenton Gregory, Carmen A Puliafito, et al. Optical coherence tomography. science, 254(5035):1178–1181, 1991.

15. Luiza Larsen, Lewis D Griffin, David Gräßel, Otto W Witte, and Hubertus Axer. Polarized light imaging of white matter architecture. Microscopy research and technique, 70(10):851–863, 2007.

16. Kwanghun Chung and Karl Deisseroth. CLARITY for mapping the nervous system. Nature methods, 10(6):508, 2013.

17. Mian Wei, Lingyan Shi, Yihui Shen, Zhilun Zhao, Asja Guzman, Laura J Kaufman, Lu Wei, and Wei Min. Volumetric chemical imaging by clearing-enhanced stimulated raman scattering microscopy. Proceedings of the National Academy of Sciences, 116(14):6608–6617, 2019.

18. Kunie Ando, Quentin Laborde, Jean-Pierre Brion, and Charles Duyckaerts. 3d imaging in the postmortem human brain with clarity and cubic. In Handbook of clinical neurology, volume 150, pages 303–317. Elsevier, 2018.

19. Satoshi Nojima, Etsuo A Susaki, Kyotaro Yoshida, Hiroyoshi Takemoto, Naoto Tsujimura, Shohei Iijima, Ko Takachi, Yujiro Nakahara, Shinichiro Tahara, Kenji Ohshima, et al. Cubic pathology: three-dimensional imaging for pathological diagnosis. Scientific reports, 7(1):9269, 2017.

20. Eva L Dyer, William Gray Roncal, Judy A Prasad, Hugo L Fernandes, Doga Gürsoy, Vincent De Andrade, Kamel Fezzaa, Xianghui Xiao, Joshua T Vogelstein, Chris Jacobsen, et al. Quantifying mesoscale neuroanatomy using x-ray microtomography. eneuro, 4(5), 2017.

21. Lamiae Abdeladim, Katherine S Matho, Solène Clavreul, Pierre Mahou, Jean-Marc Sintes, Xavier Solinas, Ignacio Arganda-Carreras, Stephen G Turney, Jeff W Lichtman, Anatole Chessel, et al. Multicolor multiscale brain imaging with chromatic multiphoton serial microscopy. Nature communications, 10(1):1662, 2019.

22. HuBMAP Consortium. The human body at cellular resolution: the nih human biomolecular atlas program. Nature, 574(7777):187, 2019.

23. Colleen Bailey, Bernard Siow, Eleftheria Panagiotaki, John H Hipwell, Thomy Mertzanidou, Julie Owen, Patrycja Gazinska, Sarah E Pinder, Daniel C Alexander, and David J Hawkes. Microstructural models for diffusion mri in breast cancer and surrounding stroma: an ex vivo study. NMR in Biomedicine, 30(2):e3679, 2017.

24. Thomy Mertzanidou, John H Hipwell, Sara Reis, David J Hawkes, Babak Ehteshami Bejnordi, Mehmet Dalmis, Suzan Vreemann, Bram Platel, Jeroen van der Laak, Nico Karssemeijer, et al. 3d volume reconstruction from serial breast specimen radiographs for mapping between histology and 3d whole specimen imaging. Medical physics, 44(3):935–948, 2017.

25. Mauricio Kugler, Yushi Goto, Yuki Tamura, Naoki Kawamura, Hirokazu Kobayashi, Tatsuya Yokota, Chika Iwamoto, Kenoki Ohuchida, Makoto Hashizume, Akinobu Shimizu, et al. Robust 3d image reconstruction of pancreatic cancer tumors from histopathological images with different stains and its quantitative performance evaluation. International journal of computer assisted radiology and surgery, pages 1–9, 2019.

26. J Scott Cordova, Saumya S Gurbani, Jeffrey J Olson, Zhongxing Liang, Lee AD Cooper, Hui-Kuo G Shu, Eduard Schreibmann, Stewart G Neill, Constantinos G Hadjipanayis, Chad A Holder, et al. A systematic pipeline for the objective comparison of whole-brain spectroscopic mri with histology in biopsy specimens from grade 3 glioma. Tomography, 2(2):106, 2016.

27. Hernan Morales-Navarrete, Fabian Segovia-Miranda, Piotr Klukowski, Kirstin Meyer, Hidenori Nonaka, Giovanni Marsico, Mikhail Chernykh, Alexander Kalaidzidis, Marino Zerial, and Yannis Kalaidzidis. A versatile pipeline for the multi-scale digital reconstruction and quantitative analysis of 3d tissue architecture. Elife, 4:e11214, 2015.

28. Menuka Pallebage-Gamarallage, Sean Foxley, Ricarda AL Menke, Istvan N Huszar, Mark Jenkinson, Benjamin C Tendler, Chaoyue Wang, Saad Jbabdi, Martin R Turner, Karla L Miller, et al. Dissecting the pathobiology of altered mri signal in amyotrophic lateral sclerosis: A post mortem whole brain sampling strategy for the integration of ultra-high-field mri and quantitative neuropathology. BMC neuroscience, 19(1):11, 2018.

29. Daniel H Adler, Laura EM Wisse, Ranjit Ittyerah, John B Pluta, Song-Lin Ding, Long Xie, Jiancong Wang, Salmon Kadivar, John L Robinson, Theresa Schuck, et al. Characterizing the human hippocampus in aging and alzheimer’s disease using a computational atlas derived from ex vivo mri and histology. Proceedings of the National Academy of Sciences, 115(16):4252–4257, 2018.

30. Juan Eugenio Iglesias, Ricardo Insausti, Garikoitz Lerma-Usabiaga, Martina Bocchetta, Koen Van Leemput, Douglas N Greve, Andre Van der Kouwe, Bruce Fischl, César Caballero-Gaudes, Pedro M Paz-Alonso, et al. A probabilistic atlas of the human thalamic nuclei combining ex vivo mri and histology. Neuroimage, 183:314–326, 2018.

31. Marcel Weiss, Anneke Alkemade, Max C Keuken, Christa Műller-Axt, Stefan Geyer, Robert Turner, and Birte U Forstmann. Spatial normalization of ultrahigh resolution 7 t magnetic resonance imaging data of the postmortem human subthalamic nucleus: a multistage approach. Brain Structure and Function, 220(3):1695–1703, 2015.

32. Maged Goubran, Cathie Crukley, Sandrine de Ribaupierre, Terence M Peters, and Ali R Khan. Image registration of ex-vivo mri to sparsely sectioned histology of hippocampal and neocortical temporal lobe specimens. Neuroimage, 83:770–781, 2013.

33. Daniel H Adler, John Pluta, Salmon Kadivar, Caryne Craige, James C Gee, Brian B Avants, and Paul A Yushkevich. Histology-derived volumetric annotation of the human hippocampal subfields in postmortem mri. Neuroimage, 84:505–523, 2014.

34. Samuel CD Cartmell, Qiyuan Tian, Brandon J Thio, Christoph Leuze, Li Ye, Nolan R Williams, Grant Yang, Gabriel Ben-Dor, Karl Deisseroth, Warren M Grill, et al. Multimodal characterization of the human nucleus accumbens. NeuroImage, 198:137–149, 2019.

35. Ana Tereza Di Lorenzo Alho, C Hamani, EJL Alho, RE da Silva, GAB Santos, RC Neves, LL Carreira, CM. Araújo, G Magalhães, DB Coelho, et al. Magnetic resonance diffusion tensor imaging for the pedunculopontine nucleus: proof of concept and histological correlation. Brain Structure and Function, 222(6):2547–2558, 2017.

36. Roger M Bourne, Colleen Bailey, Edward William Johnston, Hayley Pye, Susan Heavey, Hayley Whitaker, Bernard Siow, Alex Freeman, Greg L Shaw, Ashwin Sridhar, et al. Apparatus for histological validation of in vivo and ex vivo magnetic resonance imaging of the human prostate. Frontiers in oncology, 7:47, 2017.

37. Colleen Bailey, Roger M Bourne, Bernard Siow, Edward W Johnston, Mrishta Brizmohun Appayya, Hayley Pye, Susan Heavey, Thomy Mertzanidou, Hayley Whitaker, Alex Freeman, et al. Verdict mri validation in fresh and fixed prostate specimens using patient-specific moulds for histological and mr alignment. NMR in Biomedicine, 32(5):e4073, 2019.

38. Arnauld Sergé, Anne-Laure Bailly, Michel Aurrand-Lions, Beat A Imhof, and Magali Irla. For3d: full organ reconstruction in 3d, an automatized tool for deciphering the complexity of lymphoid organs. Journal of immunological methods, 424:32–42, 2015.

39. Rushin Shojaii, Stephanie Bacopulos, Wenyi Yang, Tigran Karavardanyan, Demetri Spyropoulos, Afshin Raouf, Anne Martel, and Arun Seth. Reconstruction of 3-dimensional histology volume and its application to study mouse mammary glands. JoVE (Journal of Visualized Experiments), 89:e51325, 2014.

40. Herbert Thiele, Stefan Heldmann, Dennis Trede, Jan Strehlow, Stefan Wirtz, Wolfgang Dreher, Judith Berger, Janina Oetjen, Jan Hendrik Kobarg, Bernd Fischer, et al. 2d and 3d maldi-imaging: conceptual strategies for visualization and data mining. Biochimica et Biophysica Acta (BBA)-Proteins and Proteomics, 1844(1):117–137, 2014.

41. Michel E Vandenberghe, Anne-Sophie Hérard, Nicolas Souedet, Elmahdi Sadouni, Mathieu D Santin, Dominique Briet, Denis Carré, Jocelyne Schulz, Philippe Hantraye, Pierre-Etienne Chabrier, et al. High-throughput 3d whole-brain quantitative histopathology in rodents. Scientific reports, 6:20958, 2016.

42. HB Stolp, G Ball, P-W So, J-D Tournier, M Jones, C Thornton, and AD Edwards. Voxel-wise comparisons of cellular microstructure and diffusion-mri in mouse hippocampus using 3d bridging of optically-clear histology with neuroimaging data (3d-bond). Scientific reports, 8(1):4011, 2018.

43. Maik Stille, Edward J Smith, William R Crum, and Michel Modo. 3d reconstruction of 2d fluorescence histology images and registration with in vivo mr images: application in a rodent stroke model. Journal of neuroscience methods, 219(1):27–40, 2013.

44. Meng Kuan Lin, Yeonsook Shin Takahashi, Bing-Xing Huo, Mitsutoshi Hanada, Jaimi Nagashima, Junichi Hata, Alexander S Tolpygo, Keerthi Ram, Brian C Lee, Michael I Miller, et al. A high-throughput neurohistological pipeline for brain-wide mesoscale connectivity mapping of the common marmoset. Elife, 8:e40042, 2019.

45. Peizhen Sun, Prasanna Parvathaneni, Kurt G Schilling, Yurui Gao, Vaibhav Janve, Adam Anderson, and Bennett A Landman. Integrating histology and mri in the first digital brain of common squirrel monkey, saimiri sciureus. In Medical Imaging 2015: Biomedical Applications in Molecular, Structural, and Functional Imaging, volume 9417, page 94171T. International Society for Optics and Photonics, 2015.

46. D Baldi, M Aiello, A Duggento, M Salvatore, and C Cavaliere. Mr imaging-histology correlation by tailored 3d-printed slicer in oncological assessment. Contrast Media & Molecular Imaging, 2019, 2019.

47. Joseph R Guy, Pascal Sati, Emily Leibovitch, Steven Jacobson, Afonso C Silva, and Daniel S Reich. Custom fit 3d-printed brain holders for comparison of histology with mri in marmosets. Journal of neuroscience methods, 257:55–63, 2016.

48. Sethu K Boopathy Jegathambal, Kelvin Mok, David A Rudko, and Amir Shmuel. Mri based brain-specific 3d-printed model aligned to stereotactic space for registering histology to mri. In 2018 40th Annual International Conference of the IEEE Engineering in Medicine and Biology Society (EMBC), pages 802–805. IEEE, 2018.

49. Maryana Alegro, Edson Amaro-Jr, Burlen Loring, Helmut Heinsen, Eduardo Alho, Lilla Zollei, Daniela Ushizima, and Lea T Grinberg. Multimodal whole brain registration: Mri and high resolution histology. In Proceedings of the IEEE conference on computer vision and pattern recognition workshops, pages 194–202, 2016.

50. Shan Yang, Zhengyi Yang, Karin Fischer, Kai Zhong, Jörg Stadler, Frank Godenschweger, Johann Steiner, Hans-Jochen Heinze, Hans-Gert Bernstein, Bernhard Bogerts, et al. Integration of ultra-high field mri and histology for connectome based research of brain disorders. Frontiers in neuroanatomy, 7:31, 2013.

51. Juan Eugenio Iglesias, Marco Lorenzi, Sebastiano Ferraris, Loïc Peter, Marc Modat, Allison Stevens, Bruce Fischl, and Tom Vercauteren. Model-based refinement of nonlinear registrations in 3d histology reconstruction. In International Conference on Medical Image Computing and Computer-Assisted Intervention, pages 147–155. Springer, 2018.

52. M Mancini, S Crampsie, DL Thomas, Z Jaunmuktane, JL Holton, and JE Iglesias. Hierarchical joint registration of tissue blocks with soft shape constraints for large-scale histology of the human brain. In 2019 IEEE 16th International Symposium on Biomedical Imaging (ISBI 2019), pages 666–669. IEEE, 2019.

53. Juan Eugenio Iglesias, Shauna Crampsie, Catherine Strand, Mohamed Tachrount, David L Thomas, and Janice L Holton. Effect of fluorinert on the histological properties of formalin-fixed human brain tissue. Journal of Neuropathology & Experimental Neurology, 77(12):1085–1090, 2018.

54. John P Mugler III. Optimized three-dimensional fast-spin-echo mri. Journal of magnetic resonance imaging, 39(4):745–767, 2014.

55. Tim Holy. Maximally perceptually-distinct colors. https://www.mathworks.com/matlabcentral/fileexchange/29702-generate-maximally-perceptually-distinct-colors, 2011. Accessed: 09-03-2019.

56. Marc Modat, Gerard R Ridgway, Zeike A Taylor, Manja Lehmann, Josephine Barnes, David J Hawkes, Nick C Fox, and Sébastien Ourselin. Fast free-form deformation using graphics processing units. Computer methods and programs in biomedicine, 98(3):278–284, 2010.

57. Oula Puonti, Juan Eugenio Iglesias, and Koen Van Leemput. Fast and sequence-adaptive whole-brain segmentation using parametric bayesian modeling. NeuroImage, 143:235–249, 2016.

58. Bruce Fischl. Freesurfer. Neuroimage, 62(2):774–781, 2012.

59. William E Lorensen and Harvey E Cline. Marching cubes: A high resolution 3d surface construction algorithm. In ACM siggraph computer graphics, volume 21, pages 163–169. ACM, 1987.

60. Bruce Fischl, Martin I Sereno, and Anders M Dale. Cortical surface-based analysis: Ii: inflation, flattening, and a surface-based coordinate system. Neuroimage, 9(2):195–207, 1999.

61. Bruce Fischl, Arthur Liu, and Anders M Dale. Automated manifold surgery: constructing geometrically accurate and topologically correct models of the human cerebral cortex. IEEE transactions on medical imaging, 20(1):70–80, 2001.

62. Anders M Dale, Bruce Fischl, and Martin I Sereno. Cortical surface-based analysis: I. segmentation and surface reconstruction. Neuroimage, 9(2):179–194, 1999.

63. Rahul S Desikan, Florent Ségonne, Bruce Fischl, Brian T Quinn, Bradford C Dickerson, Deborah Blacker, Randy L Buckner, Anders M Dale, R Paul Maguire, Bradley T Hyman, et al. An automated labeling system for subdividing the human cerebral cortex on mri scans into gyral based regions of interest. Neuroimage, 31(3):968–980, 2006.

64. Herbert Bay, Tinne Tuytelaars, and Luc Van Gool. Surf: Speeded up robust features. In European conference on computer vision, pages 404–417. Springer, 2006.

65. Martin A Fischler and Robert C Bolles. Random sample consensus: a paradigm for model fitting with applications to image analysis and automated cartography. Communications of the ACM, 24(6):381–395, 1981.

66. Jonathan Long, Evan Shelhamer, and Trevor Darrell. Fully convolutional networks for semantic segmentation. In Proceedings of the IEEE conference on computer vision and pattern recognition, pages 3431–3440, 2015.

67. Karen Simonyan and Andrew Zisserman. Very deep convolutional networks for large-scale image recognition. arXiv preprint arXiv: 1409.1556, 2014.

68. Arthur P Dempster, Nan M Laird, and Donald B Rubin. Maximum likelihood from incomplete data via the em algorithm. Journal of the Royal Statistical Society: Series B (Methodological), 39(1):1–22, 1977.

69. William M Wells III, Paul Viola, Hideki Atsumi, Shin Nakajima, and Ron Kikinis. Multimodal volume registration by maximization of mutual information. Medical image analysis, 1(1):35–51, 1996.

70. Frederik Maes, Andre Collignon, Dirk Vandermeulen, Guy Marchal, and Paul Suetens. Multimodality image registration by maximization of mutual information. IEEE transactions on Medical Imaging, 16(2):187–198, 1997.

71. Josien PW Pluim, JB Antoine Maintz, and Max A Viergever. Mutual-information-based registration of medical images: a survey. IEEE transactions on medical imaging, 22(8):986–1004, 2003.

72. Dong C Liu and Jorge Nocedal. On the limited memory bfgs method for large scale optimization. Mathematical programming, 45(1-3):503–528, 1989.

73. Vincent Arsigny, Olivier Commowick, Xavier Pennec, and Nicholas Ayache. A log-euclidean framework for statistics on diffeomorphisms. In International Conference on Medical Image Computing and Computer-Assisted Intervention, pages 924–931. Springer, 2006.

74. Caroline Magnain, Jean C Augustinack, Martin Reuter, Christian Wachinger, Matthew P Frosch, Timothy Ragan, Taner Akkin, Van J Wedeen, David A Boas, and Bruce Fischl. Blockface histology with optical coherence tomography: a comparison with nissl staining. NeuroImage, 84:524–533, 2014.

75. Hui Wang, Taner Akkin, Caroline Magnain, Ruopeng Wang, Jay Dubb, William J Kostis, Mohammad A Yaseen, Avilash Cramer, Sava Sakadžić, and David Boas. Polarization sensitive optical coherence microscopy for brain imaging. Optics letters, 41(10):2213–2216, 2016.

76. Markus Axer, David Grässel, Melanie Kleiner, Jürgen Dammers, Timo Dickscheid, Julia Reckfort, Tim Hütz, Björn Eiben, Uwe Pietrzyk, Karl Zilles, et al. High-resolution fiber tract reconstruction in the human brain by means of three-dimensional polarized light imaging. Frontiers in neuroinformatics, 5:34, 2011.

77. Hei Ming Lai, Alan King Lun Liu, Harry Ho Man Ng, Marc H Goldfinger, Tsz Wing Chau, John DeFelice, Bension S Tilley, Wai Man Wong, Wutian Wu, and Steve M Gentleman. Next generation histology methods for three-dimensional imaging of fresh and archival human brain tissues. Nature communications, 9(1):1066, 2018.

78. Markus Morawski, Evgeniya Kirilina, Nico Scherf, Carsten Jäger, Katja Reimann, Robert Trampel, Filippos Gavriilidis, Stefan Geyer, Bernd Biedermann, Thomas Arendt, et al. Developing 3d microscopy with clarity on human brain tissue: Towards a tool for informing and validating mri-based histology. Neuroimage, 182:417–428, 2018.

79. Juan Eugenio Iglesias, Ender Konukoglu, Darko Zikic, Ben Glocker, Koen Van Leemput, and Bruce Fischl. Is synthesizing mri contrast useful for inter-modality analysis? In International Conference on Medical Image Computing and Computer-Assisted Intervention, pages 631–638. Springer, 2013.

80. Ian Goodfellow, Jean Pouget-Abadie, Mehdi Mirza, Bing Xu, David Warde-Farley, Sherjil Ozair, Aaron Courville, and Yoshua Bengio. Generative adversarial nets. In Advances in neural information processing systems, pages 2672–2680, 2014.

81. Jun-Yan Zhu, Taesung Park, Phillip Isola, and Alexei A Efros. Unpaired image-to-image translation using cycle-consistent adversarial networks. In Proceedings of the IEEE international conference on computer vision, pages 2223–2232, 2017.

82. Yuankai Huo, Zhoubing Xu, Hyeonsoo Moon, Shunxing Bao, Albert Assad, Tamara K Moyo, Michael R Savona, Richard G Abramson, and Bennett A Landman. Synseg-net: Synthetic segmentation without target modality ground truth. IEEE transactions on medical imaging, 38(4):1016–1025, 2018.

83. Andreas Hess, Rukun Hinz, Georgios A Keliris, and Philipp Boehm-Sturm. On the usage of brain atlases in neuroimaging research. Molecular Imaging and Biology, 20(5):742–749, 2018.

84. Martina Absinta, Govind Nair, Massimo Filippi, Abhik Ray-Chaudhury, Maria I Reyes-Mantilla, Carlos A Pardo, and Daniel S Reich. Postmortem magnetic resonance imaging to guide the pathologic cut: individualized, 3-dimensionally printed cutting boxes for fixed brains. Journal of Neuropathology & Experimental Neurology, 73(8):780–788, 2014.

85. Kiymet Kübra Yurt, Elfide Gizem Kivrak, Gamze Altun, Hamza Mohamed, Fathelrahman Ali, Hosam Eldeen Gasmalla, and Suleyman Kaplan. A brief update on physical and optical disector applications and sectioning-staining methods in neuroscience. Journal of chemical neuroanatomy, 93:16–29, 2018.

86. Alessandro Motta, Manuel Berning, Kevin M Boergens, Benedikt Staffler, Marcel Beining, Sahil Loomba, Philipp Hennig, Heiko Wissler, and Moritz Helmstaedter. Dense connectomic reconstruction in layer 4 of the somatosensory cortex. Science, 366(6469), 2019.

87. Maged Goubran, Christoph Leuze, Brian Hsueh, Markus Aswendt, Li Ye, Qiyuan Tian, Michelle Y Cheng, Ailey Crow, Gary K Steinberg, Jennifer A McNab, et al. Multimodal image registration and connectivity analysis for integration of connectomic data from microscopy to mri. Nature Communications, 10(1):1–17, 2019.

88. Kyle Milligan, Aishwarya Balwani, and Eva Dyer. Brain mapping at high resolutions: Challenges and opportunities. Current Opinion in Biomedical Engineering, 2019.

89. Luke J Edwards, Evgeniya Kirilina, Siawoosh Mohammadi, and Nikolaus Weiskopf. Microstructural imaging of human neocortex in vivo. Neuroimage, 182:184–206, 2018.

90. Joe Chalfoun, Michael Majurski, Tim Blattner, Kiran Bhadriraju, Walid Keyrouz, Peter Bajcsy, and Mary Brady. Mist: accurate and scalable microscopy image stitching tool with stage modeling and error minimization. Scientific reports, 7(1):1–10, 2017.

